# Quorum sensing signal autoinducer-2 inhibits sporulation of *Bacillus* by interacting with RapC and functions across species

**DOI:** 10.1101/2021.11.02.466875

**Authors:** Qin Xiong, Huihui Zhang, Xia Shu, Xiting Sun, Haichao Feng, Zhihui Xu, Ákos T. Kovács, Yunpeng Liu, Ruifu Zhang

## Abstract

Collective behavior of bacteria is regulated by quorum sensing (QS). Bacterial cells sense the density of the population and induce corresponding traits and developmental processes. Autoinducer-2 (AI-2) is a common QS signal that regulates behavior of both Gram-positive and Gram-negative bacteria. In spite of the plethora of processes described to be influenced by AI-2 in diverse Gram-negative bacteria, the AI-2-regulated processes in *Bacilli* are relatively unexplored. Previously, we demonstrated that AI-2 regulates root colonization of *Bacillus velezensis* SQR9, a well-studied plant beneficial rhizobacterium. Here, we describe a novel function for AI-2 in *B. velezensis* SQR9 related to development of dormant spores. AI-2 inhibited the initiation of spore development throught the phosphatase RapC and the DNA binding regulator ComA. Using mutant strains and protein-protein interaction studies, we demonstrate that AI-2 interacts with RapC to stimulate its binding to ComA and therefore inactive ComA. We further demonstrate that ComA is essential for Spo0A-regulated sporulation in *B. velezensis* SQR9. Finally, the AI-2 molecule could be shared cross species for inhibiting *Bacillus* sporulation. Our study revealed a novel function and regulation mechanism of AI-2 in sporulation inhibition of *Bacilli* that overall suggests sporulation to be a population-level decision process in *Bacilli* rather than just a individual cell behavior.

**Author summary:** Quorum sensing (QS) regulates many bacterial social behavior. Bacteria cells could moniter and respond cell density by sensing the self produced QS signals. While most QS signals are unique for either Gram-positive or Gram-negative bacteria, autoinducer-2 (AI-2) is a QS signal that could produced by both bacteria groups. However, knowledge of the mechanism of AI-2 affecting bacterial behavior is poorly understood. Here, we found AI-2 inhibite *Bacillus velezensis* SQR9 sporulation, a generally known bacterial individual behavior. We further revealed the mechanism of AI-2 influencing sporulation of *B. velezensis* SQR9 was dependent on RapC and ComA. AI-2 interacts with RapC to stimulate its binding to ComA and therefore inactive ComA, and then inhibited the Spo0A-regulated sporulation. Interestingly, we show *B. velezensis* SQR9 could also sense the AI-2 produced by other bacteria and reduce their own sporulation. Taken together we discovered the novel function of AI-2 in sporulation, which will expand the significance of QS signal that they regulate not only social behavior but also individual behavior of bacteria.

## Introduction

Microbes live in various habitats and are widely used in medicine, industry and agriculture. However, to survive and form stable communities with other microbes in the given niches are dependent on the abilities of the microbes to exert certain functions, such as antibiotic production, waste degradation, pathogenicity, or host beneficial functions. The behavior of bacteria in a population is under regulation by quorum sensing (QS) [1], a strategy during which bacterial cells sense the population density and express certain traits or induce differention processes. Study of QS has draw much attention because it provided target for manipulation of disease [2], industrial fermentation and agricultural production [3,4].

Quorum sensing is achieved through bacterial perception of signal molecules produced by individual cells that gradually increasing along with cell density, and either sensed by receptors displayed on the cell membrane or reimported into cells where it binds directly to regulatory proteins. QS has been reported to regulate many bacterial social behaviors, such as biofilm formation, virulence and symbiosis [5]. Bacterial QS signal includes N-acyl-homoserine lactones (AHLs), cholera autoinducer 1 (CAI-1), autoinducer-2 (AI-2), and some specific oligopeptides [6]. The AHLs controlled by LuxI/LuxR-like system are the most studied QS signals, however, these molecules have only been described in Gram-negative bacteria [5]. In Gram-positive bacteria, unique oligopeptide QS signals are detected by two-component system, such as CSP (competence stimulating peptide) sensed by ComD/ComE system in *Streptococcus pneumoniae* [7]. Besides these unique molecules specific to their respective bacterial group, AI-2 is a general QS signal that plays important roles in both Gram-positive and Gram-negative bacteria [8]. However, it is less explored whether and how the general AI-2 signal is function cross species.

AI-2 has diverse functions in regulating bacterial social behavior [9], including the regulation of biofilm formation in *Bacillus cereus* and *Staphylococcus aureus* [10,11], and virulence of *Streptococcus pyogenes* and enterohemorrhagic *Escherichia coli* [12,13]. As other QS signals, AI-2 mediated processes generally involve the production, release, reimportation and sensing of the signal [14]. AI-2 is synthesized from *S*-adenosylmethionine (SAM) by the LuxS enzyme [9] and subsequenctly released from the cell by free diffusion. The reimportation of AI-2 into bacterial cells was generally controlled by the LuxP/LuxQ two-component and *lsr* operon-like transport apparatus in a density dependent manner [5,15]. After being imported back to the cell, the intracellular AI-2 function as a signal that regulates various bacterial behaviors. In the Gram-negative bacterium, *Vibrio harveyi*, AI-2 regulated luminescence relies on phosphorylated LuxO, which in turn controls the expression of sRNA (*Qrr*) and LuxR, a transcription regulator of luminescence genes [16,17]. In contrast to the well explored regulatory pathways, the mechanism how AI-2 influences gene regulation in Gram-positive bacteria remains unclear.

Bacterial cells grow and interact with each other, which leads to various types of interactions. At high cell densities where bacteria secrete common goods that are used by any member of the whole population, bacteria engage in a social lifestyle (i.e. traits are influencing the whole population and not just single cells). QS systems have an important influence on these pathways. In contrast, the formation of extremely resistant, dormat cell types, bacterial spores can be regarded as non-social life stage that depends simply on the properties of the spore itself. It has been so far unexplored whether QS, a population-level regulatory system, also influences individual life style, like sporulation. Sporulation has been mostly studied in the *Bacillus* genus, a group of Gram-positive bacteria. The initiation of sporulation in *Bacilli* is mainly controlled by phosphorylation of the global regulator, Spo0A [18]. The phophorylation and therefore activation of Spo0A are controled by two ways, the phosphorylation cascade activates Spo0A by histidine kinases through Spo0F and Spo0B, while Spo0F is directly dephosphoryled by Rap phosphatases reducing the phosphate flow in the pathway. The activities of Rap phosphatases are repressed by corresponding signaling Phr peptides. Sporulation of *Bacilli* has been studied for decades and the regulatory pathway has been dissected to understand the influence of environmental stress conditions, including nutritional deficiency [19,20]. However, sporulation has been always recognized as an individual cell behavior in *Bacilli* rather than a social differentiation regulated by QS.

*Bacillus velezensis* SQR9 is a plant beneficial rhizobacterium, isolated from the rhizosphere of cucumber with the capability to promote plant growth [21]. Our previous study demonstrated that defect in the *luxS* gene, which codes for the synthesase of AI-2, caused impaired biofilm formation and root colonization by *B. velezensis* SQR9 [22]. In this study, we discovered a novel regulation function of AI-2 for inhibiting sporulation of *B. velezensis* SQR9. The molecular mechanism was also elucidated, revealing that AI-2 directly binds to RapC and stimulates its protein-protein interaction with ComA. The interaction of RapC and ComA inhibits the activity of the latter regulator, resulting in the inhibition of the sporulation process. Moreover, sporulation of *B. velezensis* SQR9 was inhibited by AI-2 produced by *E. coli*. This study revealed a novel regulatory mechanism by AI-2 on sporulation of *B. velezensis*.

## Results

### AI-2 down-regulates spore formation of *B. velezensis* SQR9

Our previous study demonstrated that AI-2 mutant of *B. velezensis* SQR9 (SQR9/Δ*luxS*) was impaired in biofilm formation and root colonization [22]. To explore how AI-2 influences gene expression in SQR9, RNA-seq was performed that revealed 1319 genes to be up-regulated and 1459 genes to be down-regulated in the SQR9 strain 12 h after treatment with AI-2. AI-2 treatment enriched various differentially expressed genes (DEGs) related to cell wall biogenesis and lipid glycosylation, which are major process contributing to sporulation (Fig. 1A) [23]. Moreover, genes related to sporulation in *B. velezensis* SQR9 were significantly down-regulated at the late growth stage after AI-2 addition, including expression of spore coat protein coding genes (Fig. S1). qRT-PCR was used to verify these findings, demonstrating that the transcription of sporulation genes has indeed been sharply reduced by excess AI-2 (Fig. 1B). Subsequently, evaluation of the sporulation frequency in response to AI-2 revealed that addition of 4 μM exogenous AI-2 decreased the spore level to 0.1% - 1% in comparison with the wild-type SQR9 frequency of about 8% in the absence of AI-2, while the AI-2 synthesis mutant SQR9/Δ*luxS* displayed 11% frequency (Fig. 1C). Importantly, titration of AI-2 showed that the sporulation frequency was decreased along with the increase of excess AI-2 (Fig. 1C). These results indicated that AI-2 significantly inhibited spore formation of *B. velezensis* SQR9.

**Figure 1.**
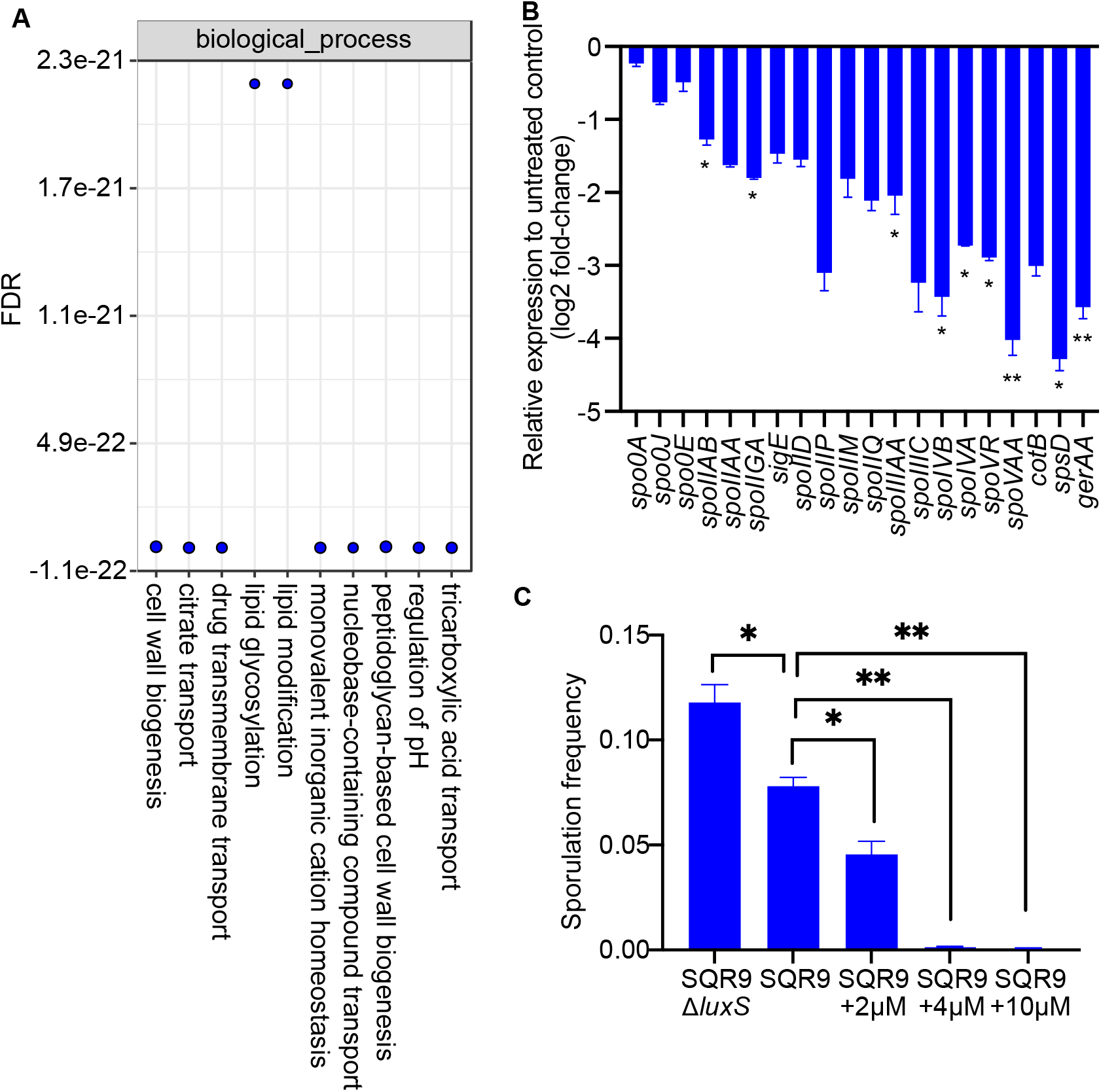
Gene expression of *B. velezensis* SQR9 induced by AI-2. (A) Gene Ontology (GO) analysis of DEGs induced by AI-2 in RNA-seq data. Most high enriched terms were primarily included for plot. Adjustment of p value was performed with BH procedure (Benjamini/Hochberg). (B) Quantitative PCR of SQR9 genes related to sporulation in samples treated with AI-2. Three independent replicates were included for each treatment and the error bars indicate the standard deviations, “*” and “**” represent significant difference (Student’s t test. p<0.05 and p<0.01, respectively) in comparison with the untreated control. (C) Sporulation frequency (the percentage of viable cells (spores) after 80 °C treatment for 20 minutes compared to the total number of untreated living cells) of *B. velezensis* SQR9, the *luxS* mutant, and AI-2 supplemented SQR9 strains cultivated in MSgg medium. Error bars indicate the standard deviations based on three independently replicated experimental values. “*” and “**” represent significant difference (Student’s t test, p<0.05 and p<0.01, respectively).

### RapC, a transcriptional regulator of Spo0A phosphatase was required for AI-2 dependent regulation of sporulation

The initiation of spore development is regulated by the global transcription factor, Spo0A in *Bacilli*. When phosphorylated Spo0A (Spo0A-P) reaches a high concentration, cells initiate formation of spores [24]. To explore how AI-2 inhibits spore formation of SQR9, the expressions of the Spo0A-P regulated genes, such as *spoIIAA*, *spoIIAB* and *spoIIGA* were investigated using qPCR that revealed down-regulation by AI-2 addition (Fig. 2A), suggesting that AI-2 might affect sporulation through the Spo0A pathway.

**Figure 2.**
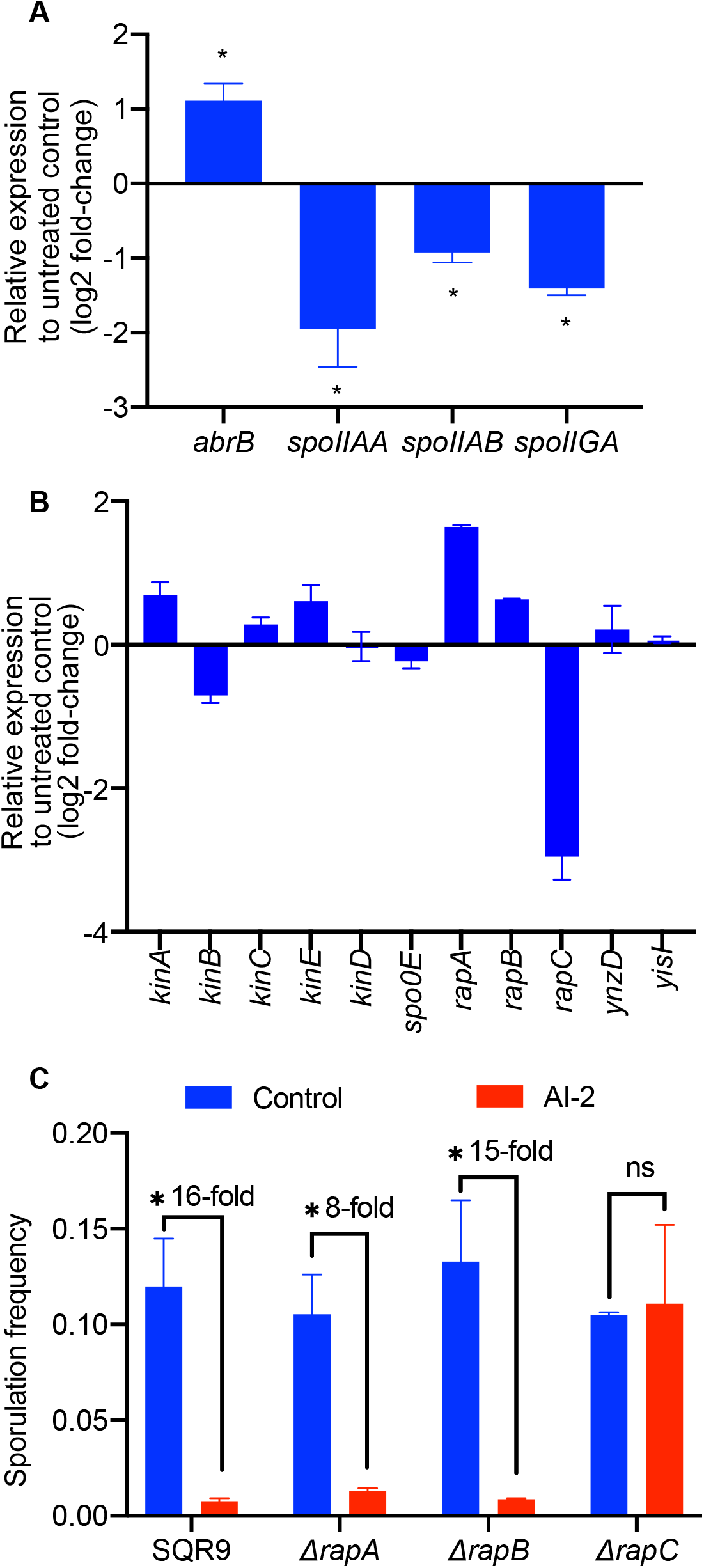
Contribution of *rapC* to sporulation inhibition by AI-2. (A) Quantitative PCR of Spo0A-P directly regulated genes in AI-2 treated samples compared with the control cultures without AI-2 addition. (B) Quantitative PCR of the genes involved in regulating phosphorylation of Spo0A. Three independent replicates were included for each treatment and the error bars indicate the standard deviations, “*” represents significant difference (p< 0.05). (C) The sporulation frequency of SQR9 and mutants in response to excess AI-2. The error bars indicate the standard deviations based on three independently replicated experimental values. “*” represents significant difference (p<0 .05), “ns” indicats no significant difference.

Spo0A phosphorylation in *Bacilli* is accomplished through a phosphate group being transfered by a cascade including Spo0F and Spo0B [25]. Spo0F is phosphorylated by one of the histidine kinases, KinA, KinB, KinC, or KinD [26], while dephosphorylated by RapA and RapB, two phosphatases that are transcriptionally regulated by RapC [27]. While KinA and KinB have been described to mainly activate sporulation in *B. subtilis*, KinC and KinD influence phosphotransfer during biofilm formation [26,28–30]. Rap phosphatases respond to cell densities via sensing their cognate Phr proteins and influence either phosphorylation or DNA-binding ability of regulators [31]. These phosphatases differentially influence among others sporulation, biofilm formation and plant colonization [32–35]. Moreover, phosphatases YnzD, YisI and Spo0E can directly dephosphorylate Spo0A-P [24]. Quantitative analysis of the expression of these kinase-coding genes (KinA, KinB, KinC and KinD) showed no significant changes upon AI-2 addition, but the genes coding for phosphatase RapA and its transcriptional regulator RapC were up- and down-regulated, respectively (Fig. 2B). To further identify which of the phosphatases or histidine kinases are involved in AI-2-dependent regulation of sporulation in *B. velezensis* SQR9, the genes coding for the respective kinases (SQR9/Δ*kinA*Δ*kinB*, SQR9/Δ*kinC*Δ*kinD*) or phosphatases (SQR9/Δ*rapA*, SQR9/Δ*rapB* and SQR9/Δ*rapC*) were deleted, and their sporulation efficiency was assayed in response to AI-2. As expected from the observations in *B. subtilis*, spore formation of kinase mutant SQR9/Δ*kinA*Δ*kinB* was significantly reduced, while the mutant SQR9/Δ*kinC*Δ*kinD* was similar to that of wild-type in *B. velezensis* (Fig. S2) [28]. However, exogenous AI-2 could still reduce the initiation of sporulation in SQR9/Δ*kinA*Δ*kinB* and SQR9/Δ*kinC*Δ*kinD* strains (Fig. S2), suggesting that the kinases are upstream of the AI-2-dependent regulation of spore development in *B. velezensis* SQR9. While the SQR9/Δ*rapA*, SQR9/Δ*rapB* and SQR9/Δ*rapC* mutants displayed comparable sporulation efficiency to the wild-type SQR9, and exogenous AI-2 could reduce sporulation in SQR9/Δ*rapA* and SQR9/Δ*rapB* mutants, deletion of *rapC* gene has completely prevented the sporulation inhibition by AI-2 (Fig. 2C), indicating that RapC is necessary for AI-2-mediated regulation of spore development in *B. velezensis* SQR9.

### ComA, regulator of *rapC* gene expression is involved AI-2-mediated sporulation inhibition

ComA is a global regulator identified in most *Bacilli* and activates gene expression of *rapC*, *phrC*, *srfA*, and *rapA* [36–38]. RapC additionally functions as a suppressor of ComA by direct binding, during which it suppresses the binding of ComA to its target promoter, including that of *rapC* [39]. This feedback mechanism leads to a balance between RapC expression and ComA activity. Interestingly, deletion of *comA* dramatically reduced sporulation of *B. velezensis* SQR9 (Fig. 3A), indicating that ComA plays an important role in initiation of sporulation. Obviously, AI-2 had no influence on sporulation in the absence of *comA*, i.e. in the absence of strong sporulation (Fig. 3A). To figure out whether AI-2 acts upstream of ComA, a known ComA target gene, *srfA* involved in the the sysnthesis of surfactin was evaluated for its transcription in the presence or absence of AI-2 using the different mutant strains. While AI-2 induced expression of *srfA* in wild-type SQR9, deletion of *rapC* or *comA* diverted the influence of AI-2 (Fig. 3B). The activity of RapC to supress ComA has been previously described to be inhibited by the PhrC pentapeptide (ERGMT), also known as the competence and sporulation-stimulating factor (CSF), which interacts with RapC to suppress its binding with ComA [37–40]. Unexpectedly, deletion of *phrC* did not influenced the AI-2-dependent inhibition of sporulation (Fig. 3B). Since ComA plays an critical role in sporulation (Fig. 3A), we speculated that AI-2 might influence the interaction between RapC and ComA to influence the sporulation, and additionaly also the transcription of *rapC*.

**Figure 3.**
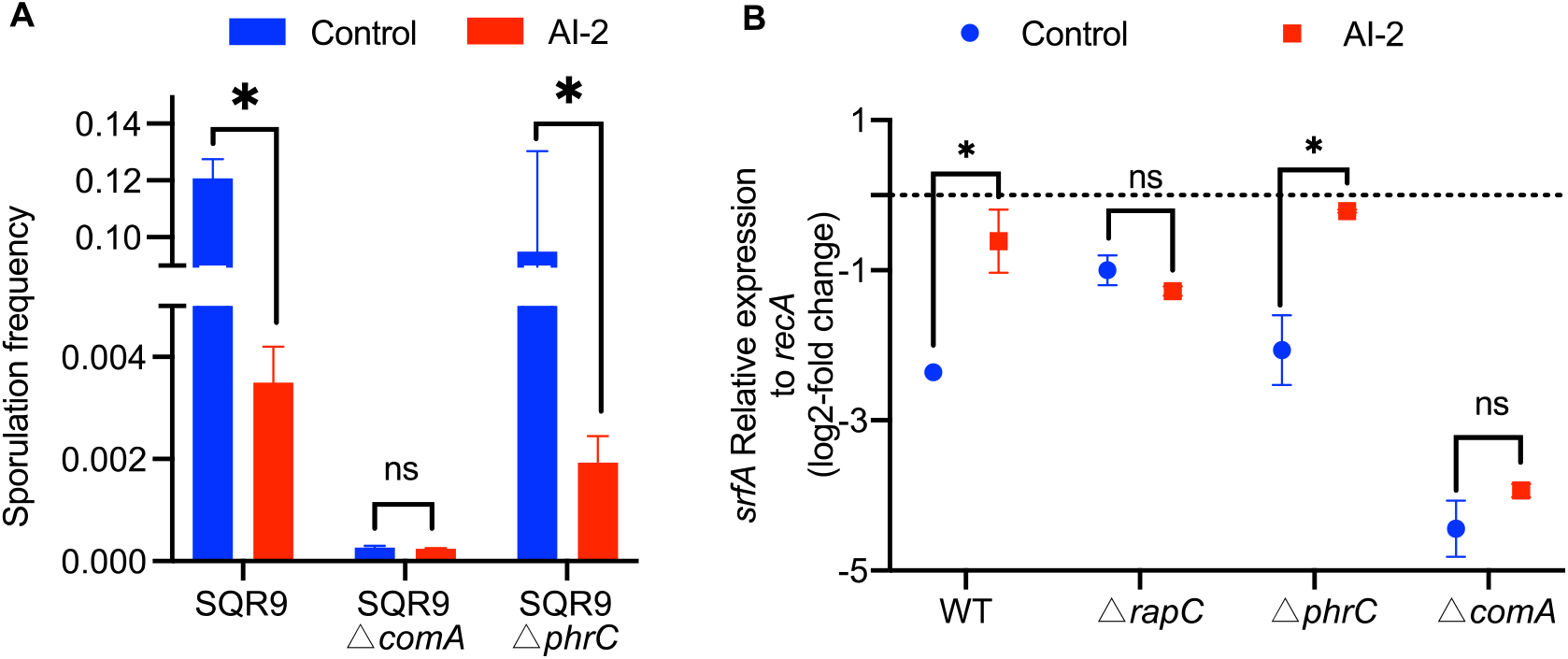
Influence of ComA on AI-2-dependent sporulation inhibition. (A) The sporulation frequency of SQR9 and mutant strains in response to excess AI-2. The error bars indicate standard deviation based on three independently replicated experimental values. “*” represents significant difference (p<0.05), “ns” indicats lack of significant difference. (B) Relative expression of *srfA* normalized to *recA* in wild-type and mutant strains with or without supplementation of AI-2. Error bars indicate standard deviation based on three independently replicated experimental values, including expression of *recA* as reference. “*” represents significant difference (p<0.05), “ns” indicats no significant difference.

### AI-2 interacts with RapC to stimulate its binding to ComA

As RapC acts as a direct repressor of ComA protein, we wondered how AI-2 influences interaction between RapC and ComA. Bio-Layer Interferometry (BLI) assay was performed to measure the binding activity between RapC and ComA, which revealed a K_D_ value of 13.5 nM between RapC and ComA (Fig. 4A). When the suppressor peptide, CSF was added, the K_D_ value for RapC-ComA binding has increased to 26.8 nM (Fig. 4A), indicating a weaker binding activity in the presence of CSF. In contrast, when AI-2 was supplemented in the assay, the K_D_ value for RapC-ComA binding has decreased to 5.49 nM, and even in the presence of CSF, AI-2 reduced the K_D_ value to 8.69 nM (Fig. 4A). These results indicate that AI-2 stimulates binding of RapC to ComA even when the respressor CSF was present. Native gel binding assays was additionally performed to evulate the direct binding of RapC to ComA that was slightly influenced by the presence of CSF (Fig. 4B). When AI-2 was added, the fraction of free RapC (i.e. unbound by ComA) was reduced, indicating that AI-2 has stimulated the binding between ComA and RapC (Fig. 4C). Collectively, AI-2 possibly inhibits sporulation in *B. velezensis* SQR9 by stimulating the binding of RapC to ComA, resulting the suppression of ComA activity.

**Figure 4.**
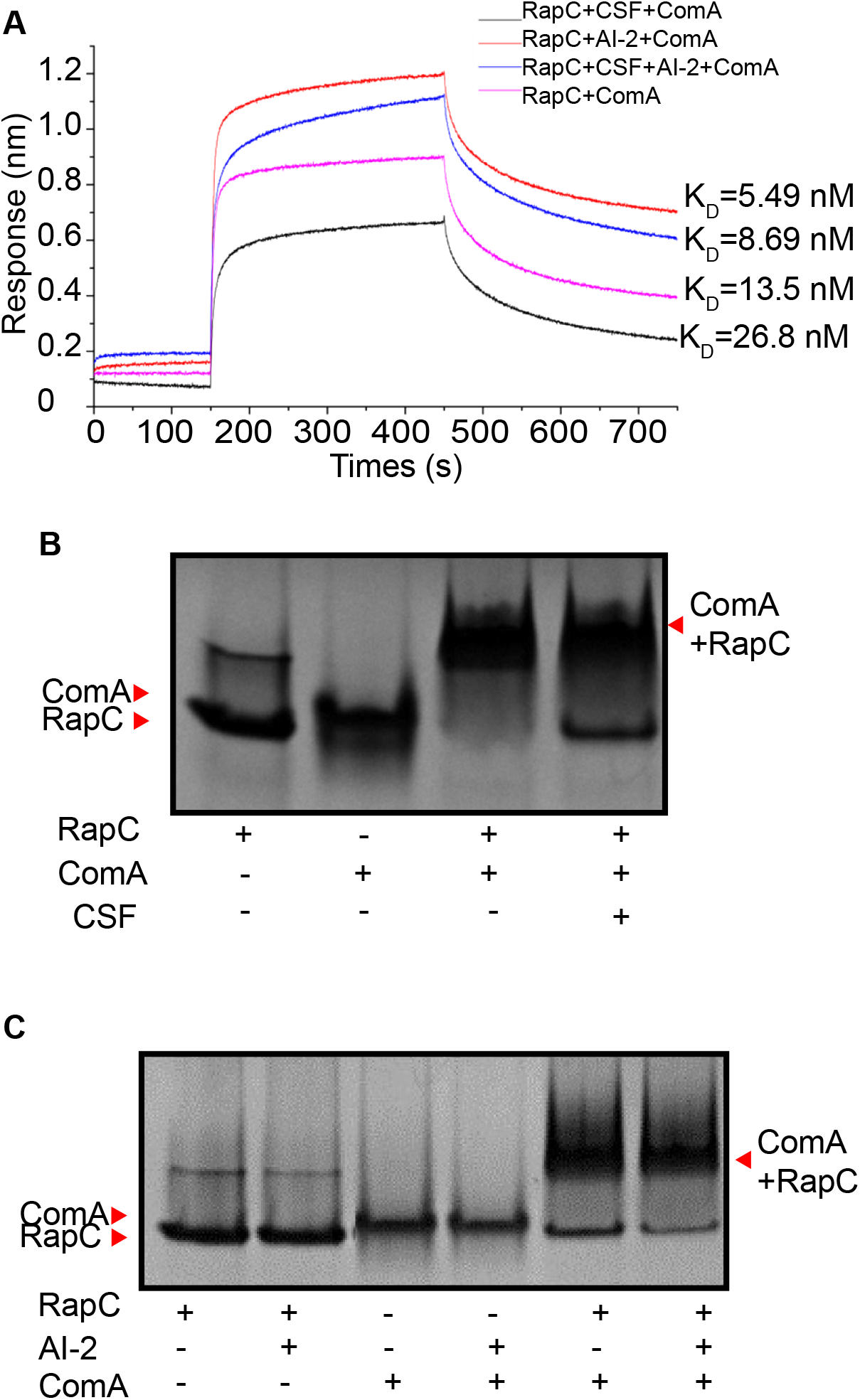
Interactions among RapC, ComA, AI-2 and CSF. (A) BLI analysis of binding activity between RapC and ComA. Super streptavidin (SSA) biosensors were loaded with RapC and two associations were subsequently performed. The first association was in solution of CSF, the second asssociation was in solution of AI-2, buffer control in each association was included in parallel experiment. Afterwards, the biosensor was associated with ComA. Buffer was included in each step in parallel for deduction in final calculation. K_D_ was recored as binding activity. (B) Native gel binding assay between RapC and ComA in the absence or presence of CSF. (C) Native gel binding assay between RapC and ComA in the absence or presence of AI-2.

### The AI-2 molecule is a cross-species signals that influence sporulation

Since structure of AI-2 is conserved among different species, we wondered whether the AI-2 signal produced by a Gram-negative species could influence the sporulation of a Gram-positive microorganism, *B. velezensis*. Therefore, an AI-2 producing bacterium, *Escherichia coli* BL21 was exploited to investigate its effect on sporulation of *B. velezensis* SQR9. After treating *B. velezensis* SQR9 with the fermentation supernatant of *E. coli* BL21, as expected, sporulation efficiency of *B. velezensis* SQR9 was significantly reduced (Fig. 5). However, when AI-2 synthesis was blocked in *E. coli* BL21 by disrupting the *luxS* gene, the fermentation supernatant failed to inhibit sporulation of *B. velezensis* SQR9 (Fig. 5). These results indicate AI-2 is not only a self-density sensing molecule, but is also involved in density sensing cross species to affect the life style of *Bacilli*.

**Figure 5.**
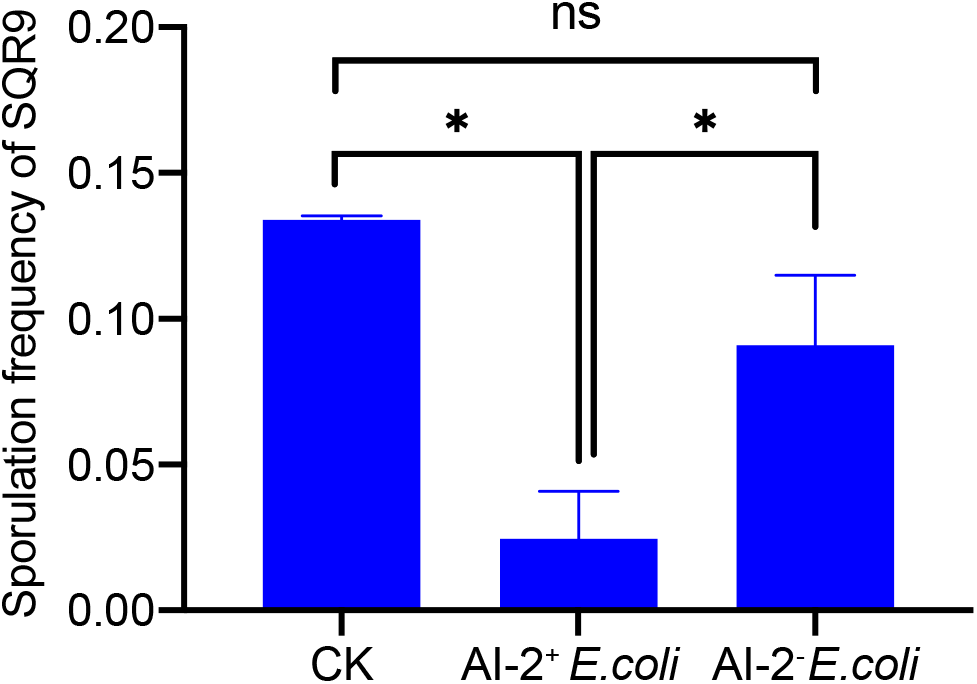
Effect of heterologous AI-2 on sporulation of *B. velezensis* SQR9. Sporulation frequency of SQR9 in response to supernatant of wild-type *E. coli* BL21 (AI-2^+^) and that of *luxS* mutant (AI-2^−^). The error bars indicate standard deviation based on three independently replicated experimental values. “*” represents significant difference (p<0 .05), “ns” indicats lack of significant difference.

## Discussion

AI-2 is an important QS signal regulating various bacterial behaviors in both Gram-negative and Gram-positive bacteria, but the regulation mechanism in Gram-positive bacteria remains unclear. Additionally, AI-2 has never been reported to inhibit sporulation of *Bacilli*. In this study, we revealed that AI-2 inhibits sporulation of *B. velezensis*, associating a novel functionality to AI-2. AI-2 seems to directly interact with RapC by stimulating its repression of ComA activity, a positive regulator of sporulation, and therefore inhibiting sporulation. This novel regulatory mechansism by AI-2 on reducing initiation of sporulation is summarized in Fig. 6. Our study additionally revealed a cross-species regulatory function of AI-2 from the Gram-negative *E. coli* to Gram-positive *B. velezensis*.

**Figure 6.**
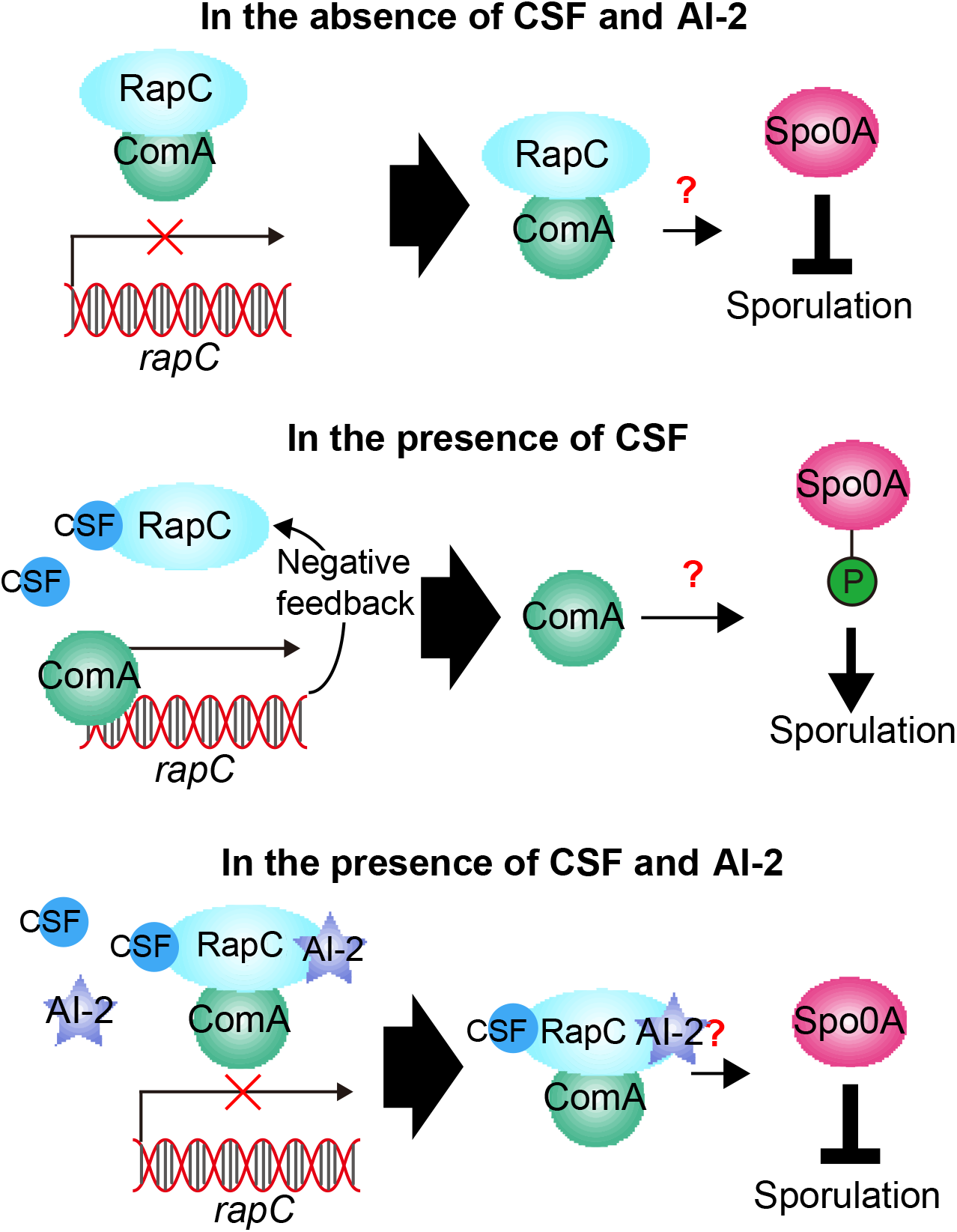
Shcematic representation of AI-2 influence on sporulation. Initiation of *B. velezensis* sporulation depends on the level of Spo0A phosphorylation, which is regulated by ComA via an unknown mechanism. RapC represses the DNA-binding ability of ComA by direct protein-protein interaction. In presence of CSF, the binding of RapC with ComA is suppressed, while in the presence of AI-2, the binding of RapC with ComA is enhanced, leading to the reduced phosphorylation of Spo0A and diminished initiation of sporulation.

AI-2 is known to regulate numerous bacterial behaviors, including motility, virulence, biofilm formation, bioluminescence, colony morphology and the type III secretion [9]. For example, AI-2 regulates motility of *Helicobacter pylori*, *E. coli* and *Campylobacter jejuni* [41,13,42], production of virulence factors in *Pseudomonas aeruginosa*, *Vibrio cholerae* and *Streptococcus pyogenes* [43,44,12], and also the expression of bioluminescence, biofilm formation, type III secretion, and protease production in *V. harveyi* [16]. Cell-cell communication signals (e.g. Phr peptides) have so far been suggested to positively influence initiation of sporulation. In contrast, AI-2 seems to inhibit the initiation of sporulation. It could be speculated that in addition to inhibition or at least delay of bacterial dormancy by AI-2, it might positively enhance plant beneficial effects, biocontrol abilities, and formation of biofilm formation as previous demonstrated [22]. Our study thus expanded the mechnisms influenced by QS signals acting in bacterial communities.

*Bacilli* are well-known model microbes for sporulation, a well-studied dormant process initiated by sensing of disadvantageous environmental cues such as nutrient exhaustion [45,46]. As spore formation in *Bacilli* eventuates at single cell level, it is regarded as an individual differentiation process rather than being a social trait. In contrast, QS is a typical social behavior that bacteria detect the cell density and exert function as a population [4]. Our results in this study indicated that sporulation is not only influenced by individual bacterial behavior but also by the population-level QS process. The observation that *E. coli* was capable of inhibiting sporulation of *B. velezensis* via AI-2 highlights that initiation of sporulation might be influenced at bacterial community level, not simply within the species or genus. At bacterial community level, delaying sporulation in the presence of high AI-2 concentration might create a critical “decision” by the *Bacillus* cells to avoid falling into dormancy even at low nutrient availability and to eliminate competitors that produce AI-2 to acquire the potentially available nutrients. Indeed, *Bacilli* produce a plethora of secondary metabolites that able to kill and lyse other bacterial cells [47]. However, this speculation needs further research in the furture. Notably, *B. subtilis* has been described to delay sporulation by lysing the cells in the population that did not activiated the Spo0A pathway, called cannibalism in *Bacilli* [48].

Quorum sensing has been studied for a long time, but most studies were aimed at Gram-negative bacteria and its major QS molecule, AHLs [49]. We previous found that AI-2 regulates biofilm formation of *B. velezensis* SQR9 [22], while AI-2 has also been reported to regulate many other physiological process of bacteria [9], however, the molecular mechnisms remain mostly unkown in *Bacilli*. Three cell-cell signaling systems are known in the *B. subtilis* group, two are regulated by oligopeptides, ComX [50] and CSF, while AI-2 is a general signaling molecule possibly recognized by most bacteria producing this substance [9]. Interestingly, all of these three molecules exert direct or indirect influence as cell-cell signaling molecule by modulating the activity of ComA [51]. CSF enhances ComA activity and thereby stimulates downstream gene expression by binding to RapC, the repressor of ComA [52]. While the transcription activatory function of ComA also relies on its phosphorylation, which is regulated by the signaling peptide ComX via the membrane-bound histidine kinase ComP [50], the repression of ComA by RapC is independent with the phosphorylation of ComA [52]. In this study, we discovered that the negative effect of AI-2 on the transcription activation of ComA is dependent on both RapC and ComA, but not on CSF. Althought both CSF and AI-2 influence RapC, their effect is opposite. Our proposed model in *B. velezensis* depicts that ComA-repressor activity and the expression of RapC creates a negative feedback loop, in which RapC repress ComA to reduce the activation of *rapC* expression by ComA. When CSF is present, the balance is shifted, as repressory role of ComA is reduced, while if AI-2 is present, repression by ComA is enhanced (Fig. 6). This effect could also explain the reduced expression of *rapC* when AI-2 is added (Fig. 2B). The AI-2 mode of action in the Gram-positive *B. velezensis* is very different from that of the Gram-negative *Vibrio* species. In *Vibrio*, AI-2 binds to LuxPQ, a membrane-bound histine kinase acting as a receptor, to regulate the phosphorylation of LuxO, the response regulator that act as repressor of downstream gene expression when it’s phosphorylation level is high [53]. Here, we revealed for the first time how AI-2 overide the influence of another QS signal, CSF and therefore influence ComA regulated processes, including activation of sporulation. These results largely expand the knowledge of QS signaling in bacteria, especially in *Bacilli*. Surprisingly, we noticed that AI-2 induced *srfA* expression (Fig. 3B) and reduced sporulation (Fig. 3A) in ComA-dependent manner, but phophoralated ComA induced both synthesis of surfactin and sporulation [54]. Moreover, in the presence of AI-2, swarming of SQR9 that depends on ComA, was also increased [22]. The opposite influence of AI-2 on the synthesis of surfactin and initiation of sporulation in *Bacilli* is unexpected and future research will be need to describe the underlying mechanism. It may be hypothesized that the ComA regulon is differentially affected depending on the interaction partner of ComA, as has been observed for SinR that represses biofilm genes, however, its target specificity is altered upon SlrR binding to SinR [55], when it represses genes related to motility and autolysins. Possibly, inhibition of sporulation might allow the population to remain metabolically active and produce yet more surfactin.

Structure of AHLs are variable among species even from one strain to another, however, the structure of AI-2 is conserved in both Gram-positive and Gram-negative bacteria [56]. It suggests that AI-2 might circulate as a shared signaling molecule in microbial communities, and therefore can be produced and sensed by distinct bacteria in a certain local environment, however, this suggestion requires further confirmation in complex microbiomes. Here, we demonstrated that *E. coli* BL21 influences sporulation of *B. velezensis* and confirmed the contribution of AI-2 in this process. AI-2 can be produced by various bacteria in a community, and sensed as a public signal molecule by different bacterial members. Due to the function of AI-2 affecting chemotaxis, biofilm formation and virulence of bacteria [57], the property as an inter-species signal indicates that it might act during microbiome assembly and might contribute to the regulation of functional traits within diverse bacteria in a certain environmental niche. For example, supplementation of an AI-2 producing strain or the AI-2 molecule directly to a synthetic rhizosphere beneficial bacterial community might regulate its colonization of the plant root, sporulation, and other beneficial functions.

## Materials and Methods

### Strains and culture conditions

The strains and plasmids used in this study are shown in Table S1. *B. velezensis* strain SQR9 (CGMCC accession no.5808, China General Microbiology Culture Collection Center, NCBI accession NO. CP006890) and its derivative strains were routinely grown at 37 °C in lysogeny broth (LB) medium (1% (w/v) tryptone, 0.5% (w/v) yeast extract, and 0.5% (w/v) NaCl). For spore formation experiments, SQR9 and its derivative strains were cultivated 12 h in MSgg medium (100 mM 3-(N-morpholino) propane sulfonic acid (MOPS), 5 mM potassium phosphate, 2 mM MgCl_2_, 700 μM CaCl_2_, 50 μM MnCl_2_, 50 μM FeCl_3_, 1 μM ZnCl_2_, 2 μM thiamine, 0.5% (w/v) glycerol, 0.5% (w/v) glutamate, 50 μg/mL tryptophan, 50 μg/mL phenylalanine, and 50 μg/mL threonine, pH 7.0) [58]. *E. coli* BL21(DE3) Δ*luxS*, which was used to express ComA and RapC proteins, was cultured in LB at 37 °C and induced for protein expression at 16 °C with 0.5 mM IPTG. *E. coli* Top10 was used for plasmid constructions and propagation. Antibiotics were used at the following concentrations: erythromycin, 1 μg/mL; zeocin, 20 μg/mL and spectinomycin, 100 μg/mL for *B. velezensis* strains, ampicillin 100 mg/mL and kanamycin 30 mg/mL for *E. coli*.

### Mutant construction

The *rapA* and *rapC* deficient marker-free mutants of *B. velezensis* were constructed using the *Pbc*-*pheS*\*-*cat* (PC) cassette and overlap-PCR based strategy as described by Xu et *al* [59]. The *kinA* and *kinB*, *kinC* and *kinD* double mutants were constructed using the same method. All the mutants were verified by DNA sequencing.

### Protein expression and purification

The plasmid pCold TF DNA was used for expressing RapC. The recombined plasmid carrying RapC coding sequence was transformed into *E. coli* BL21 (DE3)Δ*luxS* for expression. The transformed *E. coli* BL21 (DE3)Δ*luxS* was cultured at 37 °C to OD_600_ of 0.5. Isopropyl-β-D-thiogalactopryanoside (IPTG) was added with final concentration of 0.5 mM to induce protein expression at 16 °C overnight. The protein was purified from the lysate of the bacterial cells by NGC Chromatography system (BioRad, CA, USA) with nickel column. Thrombin was used to remove the tag, and nickel column was used to reverse purify the tag-free RapC protein.

The plasmid pET29a(+) was used for the expression of ComA follow the method described by Liu et *al* [60]. The recombinant plasmid was transformed into the competent state of *E. coli* Top10 and *E. coli* BL21 (DE3)Δ*luxS*, the transformants were selected on LB agar medium with Kanamycin and verified by sequencing. The 6 His-tagged protein was purified by His-affinity resin chromatography. The purified proteins were collected and stored in PBS buffer at −80°C. All proteins was evaluated by Nanodrop 2000 (Thermo scientific, MA, USA).

### Quantitative and global transcription analysis

*B. velezensis* SQR9 was cultured overnight in LB medium until OD_600_ reached 1.0, and was transferred to MSgg medium with 1 % (v/v) inoculation. AI-2 was added to the medium to a final concentration of 4 μM. After 12 hours, bacterial samples were collected for RNA extraction. Total RNA extraction was performed using a Bacterial RNA kit (OMEGA, Biotek, USA) following the instruction. The extracted RNA was checked using Nanodrop 2000 (Thermo scientific, MA, USA).

For qRT-PCR, extracted RNA was reverse-transcribed into cDNA using the PrimeScript™ RT reagent Kit with gDNA Eraser (TaKaRa, Dalian, China). The gene quantitative PCR was performed using TB green Premix EX Taq (Takara) with a QuantStudio 6 Flex (Applied Biosystems, CA, USA). The following PCR program was used: cDNA was denatured for 30 s at 95 °C, followed by 40 cycles consisting of 5 s at 95 °C and 34 s at 60 °C. Primers were provided in Table S2. The 2^−ΔΔCT^ method was used to analyze the real-time PCR data. The *recA* gene was included as an internal control. Each treatment included three independent replicates. For RNA-seq, the library was prepared and sequenced on an Illumina HiSeq 4000 and 150 bp paired-end reads were generated at the Beijing Allwegene Technology Company, Beijing, China. Clean reads were obtained after quality control and mapped to the reference genome using Tophat2. RNA-Seq data was normalized to FPKM (fragments per kilobase of exon per million fragments mapped). P value was calculated using a negative binomial distribution-based test and FDR (P adjust) was calculated using BH (Benjamini/ Hochberg). Sequence data was deposited on Sequence Read Archive (SRA). The SRA accession number is PRJNA673673. Three replicates were included for each treatment.

### Determination of spore formation of *Bacillus*

The sporulation frequency of SQR9 and the derivated strains were determined at 12 h post inoculation (the time point shown differentially expressing profile of sporulation genes). Strain SQR9 was cultured in LB medium at 37 °C until OD_600_ reached to 1.0, the cultures were then inoculated into flask with 100 mL MSgg medium to a final concentration of 1% inoculum volume and cultured for 12 h at 37 °C. Afterthen, 1mL of bacterial culture was collected. One half of the culture was diluted and spread on LB agar for counting all living cells, another half was treated with 80 °C water bath for 20 min to kill the non-sporulating cells, diluted, and spread on LB agar for counting the viable spore number. The ratio of spores to the total cells was recorded as sporulation frequency. Three independent replicates were included for each treatment.

### Bio-Layer Interferometry (BLI) assays

BLI experiments were performed using an Octet RED96 instrument (Pall ForteBio, CA, USA) at 25°C in Modified Kinetics buffer (1 x PBS, 0.05 % Tween-20). Super Streptavidin (SSA) biosensors (Pall ForteBio, CA, USA) were pre-equilibrated in buffer for 10 min at room temperature. The sensor loading by RapC was performed for 5 min and the RapC concentration was 50 μg/mL. In this experiment, two associations were performed. The first association was solution I, which was a different quorum sensing signal or buffer (CSF; AI-2; (CSF+AI-2); Buffer); the second asssociation was solution II, which was a mixture of the corresponding solution I and protein ComA ((CSF+ComA); (AI-2+ComA); (CSF+AI-2+ComA); ComA); and the final dissociation was also carried out in the same solution I as the first association. The concentration of CSF and AI-2 was the same in solution I and II, and the concentration of ComA was the same in solution II. The data were analysed using the global fitting algorithm included in the Octet Data Analysis Software 9.0 (Pall ForteBio, CA, USA).

### Native polyacrylamide gel electrophoresis

The HEPES native gel electrophoresis was performed using the Precast-GL gel with 12 % polyacrylamide (Art. No. C601102, Sangon Biotech, Shanghai, China). The electrophoresis conditions were carried out according to the product instructions. All these samples were incubated in PBS. After the end of incubation, loading buffer (12 mM Tris-HCl, 0.02 % bromophenol blue, 5 % glycerol and 2.88 mM β-mercaptoethanol) was added and mixed, and samples loading volume was 20 μL.

### Treatment of SQR9 with *E. coli* culture supernatant

The wild type *E.coli* BL21 and *luxS* gene mutant (BL21(DE3)Δ*luxS*) were cultured in LB medium until OD_600_ reached 1.0. The cultures were inoculated into 100 mL MSgg medium with 1% inoculum. After 18 hours at 37 °C, the supernatant was collected by centrifugation with 10,000 rpm for 15 minutes at 4 °C. The supernatant was filtered by 0.22 μm filter and freeze-dried into powder. The powder generated from 100 mL fermentation was dissolved with 2 mL PBS, and the solution was filtered with 0.22 μm filter. Finally, 2 mL of filtrate were added to 100 mL MSgg medium, 2 mL PBS was added as control in parellel. Meanwhile, the medium was inoculated with SQR9. After culturing at 37 °C for 12 h, the sporulation frequency was measured.

## Acknowledgements

We thank Prof. Xihui Shen of College of Life Sciences, Northwest A&F University, for providing the *E. coli* BL21Δ*luxS* strain.

## Author contributions

Conceptualization, Y.L. and R.Z.; Formal Analysis, Q.X. and Y.L.; Investigation, Q.X., H.Z., X.Shu and X.Sun; Verification, H.F. and Z.X.; Writing -Original Draft, Q.X. and Y.L.; Writing - Review & Editing, Á.T.K. and R.Z.; Visualization, Y.L.; Supervision, Z.X., Á.T.K. and R.Z.; Project Administration, Y.L. and R.Z..; Funding Acquisition, R.Z..

## References

1. Silpe JE, Bassler BL. A host-produced quorum-sensing autoinducer controls a phage lysis-lysogeny decision. Cell. 2019; 176: 268–280. https://doi:10.1016/j.cell.2018.10.059PMID: 30554875

2. Defoirdt T. Quorum-sensing systems as targets for antivirulence therapy. Trends Microbiol. 2018; 26: 313–328. https://doi:10.1016/j.tim.2017.10.005 PMID: 29132819

3. Tran LSP, Nagai T, Itoh Y. Divergent structure of the ComQXPA quorum-sensing components: Molecular basis of strain-specific communication mechanism in *Bacillus subtilis*. Mol Microbiol. 2000; 37: 1159–1171. https://doi:10.1046/j.1365-2958.2000.02069.x PMID: 10972833

4. Loh J, Pierson EA, Pierson LS, Stacey G, Chatterjee A. Quorum sensing in plant-associated bacteria. Curr Opin Plant Biol. 2002; 5: 285–290. https://doi:10.1016/S1369-5266(02)00274-1PMID: 12179960

5. Miller MB, Bassler BL. Quorum sensing in bacteria. Annu Rev Microbiol. 2001; 55: 165–199. https://doi:10.1146/annurev.micro.55.1.165 PMID: 11544353

6. Wai-Leung NG, Bassler B. Bacterial quorum-sensing network architectures. Annu Rev Genet. 2009; 43: 197–222. https://doi:10.1146/annurev-genet-102108-134304.Bacterial PMID: 19686078

7. Pestova E V., Håvarstein LS, Morrison DA. Regulation of competence for genetic transformation in *Streptococcus pneumoniae* by an auto-induced peptide pheromone and a two-component regulatory system. Mol Microbiol. 1996; 21: 853–862. https://doi:10.1046/j.1365-2958.1996.501417.x PMID: 8878046

8. De Keersmaecker SCJ, Sonck K, Vanderleyden J. Let LuxS speak up in AI-2 signaling. Trends Microbiol. 2006; 14: 114–119. https://doi:10.1016/j.tim.2006.01.003 PMID: 16459080

9. Pereira CS, Thompson JA, Xavier KB. AI-2-mediated signalling in bacteria. FEMS Microbiol Rev. 2013; 37: 156–181. https://doi:10.1111/j.1574-6976.2012.00345.x PMID: 22712853

10. Auger S, Krin E, Aymerich S, Gohar M. Autoinducer 2 affects biofilm formation by *Bacillus cereus*. Appl Environ Microbiol. 2006; 72: 937–941. https://doi:10.1128/AEM.72.1.937 PMID: 16391139

11. Yu D, Zhao L, Xue T, Sun B. *Staphylococcus aureus* autoinducer-2 quorum sensing decreases biofilm formation in an *icaR*-dependent manner. BMC Microbiol. 2012; 12: 1. https://doi:10.1186/1471-2180-12-288 PMID: 23216979

12. Lyon WR, Madden JC, Levin JC, Stein JL, Caparon MG. Mutation of *luxS* affects growth and virulence factor expression in *Streptococcus pyogenes*. Mol Microbiol. 2001; 42: 145–157. https://doi:10.1046/j.1365-2958.2001.02616.x PMID: 11679074

13. Bansal T, Jesudhasan P, Pillai S, Wood TK, Jayaraman A. Temporal regulation of enterohemorrhagic *Escherichia coli* virulence mediated by autoinducer-2. Appl Microbiol Biotechnol. 2008; 78: 811–819. https://doi:10.1007/s00253-008-1359-8 PMID: 18256823

14. Eickhoff MJ, Bassler BL. SnapShot: Bacterial quorum sensing. Cell. 2018; 174: 1328–1329. https://doi:10.1016/j.cell.2018.08.003 PMID: 30142348

15. Taga ME, Miller ST, Bassler BL. Lsr-mediated transport and processing of Al-2 in *Salmonella typhimurium*. Mol Microbiol. 2003; 50: 1411–1427. https://doi:10.1046/j.1365-2958.2003.03781.x PMID: 14622426

16. Waters CM, Bassler BL. The *Vibrio harveyi* quorum-sensing system uses shared regulatory components to discriminate between multiple autoinducers. Genes Dev. 2006; 20: 2754–2767. https://doi:10.1101/gad.1466506 PMID: 17015436

17. Bassler BL, Wright M, Silverman MR. Sequence and function of LuxO, a negative regulator of luminescence in *Vibrio harveyi*. Mol Microbiol. 1994; 12: 403–412. https://doi:10.1111/j.1365-2958.1994.tb01029.x PMID: 8065259

18. Trach K, Burbulys D, Strauch M, Wu JJ, Dhillon N, Jonas R, et al. Control of the initiation of sporulation in *Bacillus subtilis* by a phosphorelay. Res Microbiol. 1991; 142: 815–823. https://doi:10.1016/0923-2508(91)90060-N PMID: 1664534

19. Tan I.S. and Ramamurthi K.S. Spore formation in *Bacillus subtilis*. Environ Microbiol Rep. 2014; 6: 212–225. https://doi:10.1111/1758-2229.12130 PMID: 24983526

20. Driks A. Overview: Development in bacteria: Spore formation in *Bacillus subtilis*. Cell Mol Life Sci. 2002; 59: 389–391. https://doi:10.1007/s00018-002-8430-x PMID: 11964116

21. Cao Y, Zhang Z, Ling N, Yuan Y, Zheng X, Shen B, et al. *Bacillus subtilis* SQR 9 can control Fusarium wilt in cucumber by colonizing plant roots. Biol Fertil Soils. 2011; 47: 495–506.

22. Xiong Q, Liu D, Zhang H, Dong X, Zhang G, Liu Y, et al. Quorum sensing signal autoinducer-2 promotes root colonization of *Bacillus velezensis* SQR9 by affecting biofilm formation and motility. Appl Microbiol Biotechnol. 2020; 104: 7177–7185. https://doi:10.1007/s00253-020-10713-w PMID: 32621125

23. Schujman GE, Grau R, Gramajo HC, Ornella L, De Mendoza D. De novo fatty acid synthesis is required for establishment of cell type-specific gene transcription during sporulation in *Bacillus subtilis*. Mol Microbiol. 1998; 29: 1215–1224. https://doi:10.1046/j.1365-2958.1998.01004.x PMID: 9767589

24. Piggot PJ, Hilbert DW. Sporulation of *Bacillus subtilis*. Curr Opin Microbiol. 2004; 7: 579–586. https://doi:10.1016/j.mib.2004.10.001 PMID: 15556029

25. Burbulys D, Trach KA, Hoch JA. Initiation of sporulation in *B. subtilis* is controlled by a multicomponent phosphorelay. Cell. 1991; 64: 545–552. https://doi:10.1016/0092-8674(91)90238-T PMID: 1846779

26. Quisel JD, Burkholder WF, Grossman AD. In vivo effects of sporulation kinases on mutant Spo0A proteins in *Bacillus subtilis*. J Bacteriol. 2001; 183: 6573–6578. https://doi:10.1128/JB.183.22.6573-6578.2001 PMID: 11673427

27. Jiang M, Grau R, Perego M. Differential processing of propeptide inhibitors of Rap phosphatases in *Bacillus subtilis*. J Bacteriol. 2000; 182: 303–310. https://doi:10.1128/JB.182.2.303-310.2000 PMID: 10629174

28. LeDeaux JR, Yu N, Grossman AD. Different roles for KinA, KinB, and KinC in the initiation of sporulation in *Bacillus subtilis*. J Bacteriol. 1995; 177: 861–863. https://doi:10.1128/jb.177.3.861-863.1995 PMID: 7836330

29. Arnaouteli S, Bamford NC, Stanley-Wall NR, Kovács ÁT. *Bacillus subtilis* biofilm formation and social interactions. Nat Rev Microbiol. 2021; 19: 600–614. https://doi:10.1038/s41579-021-00540-9 PMID: 33824496

30. Mhatre E, Monterrosa RG, Kovács ÁT. From environmental signals to regulators: Modulation of biofilm development in Gram-positive bacteria. J Basic Microbiol. 2014; 54: 616–632. https://doi:10.1002/jobm.201400175 PMID: 24771632

31. Neiditch MB, Capodagli GC, Prehna G, Federle MJ. Genetic and structural analyses of RRNPP intercellular peptide signaling of Gram-positive bacteria. Annu Rev Genet. 2017; 51: 311–333. https://doi:10.1146/annurev-genet-120116-023507 PMID: 28876981

32. Perego M, Glaser P, Hoch JA. Aspartyl-phosphate phosphatases deactivate the response regulator components of the sporulation signal transduction system in *Bacillus subtilis*. Mol Microbiol. 1996; 19: 1151–1157. https://doi:10.1111/j.1365-2958.1996.tb02460.x PMID: 8730857

33. Verdugo-Fuentes A, Gastélum G, Rocha J, de la Torre M. Multiple and overlapping functions of quorum sensing proteins for cell specialization in *Bacillus* species. J Bacteriol. 2020; 202(10): e00721–19. https://doi:10.1128/JB.00721-19 PMID: 32071096

34. Gallegos-Monterrosa R, Christensen MN, Barchewitz T, Koppenhöfer S, Priyadarshini B, Bálint B, et al. Impact of Rap-Phr system abundance on adaptation of *Bacillus subtilis*. Commun Biol. 2021; 4. https://doi:10.1038/s42003-021-01983-9 PMID: 33850233

35. Nordgaard M, Mortensen RMR, Kirk NK, Gallegos-Monterrosa R, Kovács ÁT. Deletion of Rap-Phr systems in *Bacillus subtilis* influences in vitro biofilm formation and plant root colonization. Microbiologyopen. 2021; 10: e1212. https://doi:10.1002/mbo3.1212 PMID: 34180604

36. Roggiani M, Dubnau D. ComA, a phosphorylated response regulator protein of *Bacillus subtilis*, binds to the promoter region of *srfA*. J Bacteriol. 1993; 175: 3182–3187. https://doi:10.1128/jb.175.10.3182-3187.1993 PMID: 8387999

37. Lazazzera BA, Kurtser IG, Mcquade RS, Grossman AD. An autoregulatory circuit affecting peptide signaling in *Bacillus subtilis*. J Bacteriol. 1999; 181: 5193–5200. https://doi:10.1128/jb.181.17.5193-5200.1999 PMID: 10464187

38. Mueller JP, Bukusoglu G, Sonenshein AL. Transcriptional regulation of *Bacillus subtilis* glucose starvation- inducible genes: Control of *gsiA* by the ComP-ComA signal transduction system. J Bacteriol. 1992; 174: 4361–4373. https://doi:10.1128/jb.174.13.4361-4373.1992 PMID: 1378051

39. Core L, Perego M. TPR-mediated interaction of RapC with ComA inhibits response regulator-DNA binding for competence development in *Bacillus subtilis*. Mol Microbiol. 2003; 49: 1509–1522. https://doi:10.1046/j.1365-2958.2003.03659.x PMID: 12950917

40. Solomon JM, Lazazzera BA, Grossman AD. Purification and characterization of an extracellular peptide factor that affects two different developmental pathways in *Bacillus subtilis*. Genes Dev. 1996; 10: 2014–2024. https://doi:10.1101/gad.10.16.2014 PMID: 8769645

41. Rader BA, Campagna SR, Semmelhack MF, Bassler BL, Guillemin K. The quorum-sensing molecule autoinducer 2 regulates motility and flagellar morphogenesis in *Helicobacter pylori*. J Bacteriol. 2007; 189: 6109–6117. https://doi:10.1128/JB.00246-07 PMID: 17586631

42. Elvers KT, Park SF. Quorum sensing in *Campylobacter jejuni*: Detection of a *luxS* encoded signalling molecule. Microbiology. 2002; 148: 1475–1481. https://doi:10.1099/00221287-148-5-1475 PMID: 11988522

43. Duan K, Dammel C, Stein J, Rabin H, Surette MG. Modulation of *Pseudomonas aeruginosa* gene expression by host microflora through interspecies communication. Mol Microbiol. 2003; 50: 1477–1491. https://doi:10.1046/j.1365-2958.2003.03803.x PMID: 14651632

44. Zhu J, Miller MB, Vance RE, Dziejman M, Bassler BL, Mekalanos JJ. Quorum-sensing regulators control virulence gene expression in *Vibrio cholerae*. Proc Natl Acad Sci U S A. 2002; 99: 3129–3134. https://doi:10.1073/pnas.052694299 PMID: 11854465

45. Grossman AD. Genetic networks controlling the initiation of sporulation and the development of genetic competence in *Bacillus subtilis*. Annu Rev Genet. 1995; 29: 477–508. https://doi:10.1146/annurev.ge.29.120195.002401 PMID: 8825484

46. Piggot PJ, Coote JG. Genetic aspects of bacterial endospore formation. Bacteriol Rev. 1976; 40: 908–962. https://doi:10.1128/mmbr.40.4.908-962.1976 PMID: 12736

47. Chen XH, Vater J, Piel J, Franke P, Scholz R, Schneider K, et al. Structural and functional characterization of three polyketide synthase gene clusters in *Bacillus amyloliquefaciens* FZB 42. J Bacteriol. 2006; 188: 4024–4036. https://doi:10.1128/JB.00052-06 PMID: 16707694

48. Stevick PT, Soule M, Ayala FJ. Cannibalism by sporulating bacteria. Science. 2003; 301: 510–513. https://doi:10.1126/science.1086462. PMID: 12817086

49. Papenfort K, Bassler BL. Quorum sensing signal-response systems in Gram-negative bacteria. Nat Rev Microbiol. 2016; 14: 576–588. https://doi:10.1038/nrmicro.2016.89 PMID: 27510864

50. Magnuson R, Solomon J, Grossman AD. Biochemical and genetic characterization of a competence pheromone from *B. subtilis*. Cell. 1994; 77: 207–216. https://doi.org/10.1016/0092-8674(94)90313-1 PMID: 8168130

51. Perego M. A peptide export-import control circuit modulating bacterial development regulates protein phosphatases of the phosphorelay. Proc Natl Acad Sci U S A. 1997; 94: 8612–8617. https://doi:10.1073/pnas.94.16.8612 PMID: 9238025

52. Pottahil M, Lazazzera BA. The extracellular Phr pepetide-Rap phosphatase signaling circuit of *Bacillus subtilis*. Peptides 2003; 8: 32–45. https://doi:10.2741/913 PMID: 12456319

53. Herzog R, Peschek N, Fröhlich KS, Schumacher K, Papenfort K. Three autoinducer molecules act in concert to control virulence gene expression in *Vibrio cholerae*. Nucleic Acids Res. 2019; 47: 3171–3183. https://doi:10.1093/nar/gky1320 PMID: 30649554

54. Liang Z, Qiao JQ, Li PP, Zhang LL, Qiao ZX, Lin L, et al. A novel Rap-Phr system in *Bacillus velezensis* NAU-B3 regulates surfactin production and sporulation via interaction with ComA. Appl Microbiol Biotechnol. 2020; 104: 10059–10074. https://doi:10.1007/s00253-020-10942-z PMID: 33043389

55. Chai Y, Kolter R, Losick R. Reversal of an epigenetic switch governing cell chaining in *Bacillus subtilis* by protein instability. Mol Microbiol. 2010; 78: 218–229. https://doi:10.1111/j.1365-2958.2010.07335.x PMID: 20923420

56. Vendeville A, Winzer K, Heurlier K, Tang CM, Hardie KR. Making “sense” of metabolism: Autoinducer-2, LuxS and pathogenic bacteria. Nat Rev Microbiol. 2005; 3: 383–396. https://doi:10.1038/nrmicro1146 PMID: 15864263

57. Xavier KB, Bassler BL. LuxS quorum sensing: more than just a numbers game. Curr Opin Microbiol. 2003; 6: 191–197. https://doi:10.1016/S1369-5274(03)00028-6 PMID: 12732311

58. Steven S. Branda, Jose′ Eduardo Gonza′ lez-Pastor, Sigal Ben-Yehuda, Richard Losick and Robert K. Fruiting body formation by *Bacillus subtilis*. Proc Natl Acad Sci U S A. 2001; 98: 11621–11626. https://doi:10.1073/pnas.191384198 PMID: 11572999

59. Xu Z, Xie J, Zhang H, Wang D, Shen Q, Zhang R. Enhanced control of plant wilt disease by a xylose-inducible *degQ* gene engineered into *Bacillus velezensis* strain SQR9XYQ. Phytopathology. 2019;109: 36–43. https://doi:10.1094/PHYTO-02-18-0048-R PMID: 29927357

60. Liu Y, Feng H, Chen L, Zhang H, Dong X, Xiong Q, et al. Root-secreted spermine binds to *Bacillus amyloliquefaciens* SQR9 histidine kinase KinD and modulates biofilm formation. Mol Plant-Microbe Interact. 2020; 33: 423–432. https://doi:10.1094/MPMI-07-19-0201-R PMID: 31741422

61. Zhou C., Shi L., Ye B., Feng H., Zhang J., Zhang R.F., et al. *pheS* *, an effective host-genotype-independent counter-selectable marker for marker-free chromosome deletion in *Bacillus amyloliquefaciens*. Appl. Microbiol. Biotechnol. 2017; 101: 217–227. https://doi:10.1007/s00253-016-7906-9 PMID: 27730334

62. Yan X., Yu H.J., Hong Q., Li S.P. Cre/*lox* system and PCR-based genome engineering in *Bacillus subtilis*. Appl. Environ. Microbiol. 2008; 74: 5556–5562. https://doi:10.1128/AEM.01156-08 PMID: 18641148

